# VIBE: An R-package for advanced RNA-seq data exploration, disease stratification and therapeutic targeting

**DOI:** 10.1101/2023.10.04.560641

**Authors:** Indu Khatri, Saskia D van Asten, Leandro F. Moreno, Brandon W Higgs, Christiaan Klijn, Francis Blokzijl, Iris CRM Kolder

## Abstract

**Background:** Development of therapies e.g. antibody-based treatments, rely on several factors, including the specificity of target expression and characterization of downstream signaling pathways. While existing tools for analyzing and visualizing RNA-seq data offer evaluation of individual gene-level expression, they lack a comprehensive assessment of pathway-guided analysis, relevant for single- and dual-targeting therapeutics. Here, we introduce VIBE (<u>VI</u>sualization of <u>B</u>ulk RNA <u>E</u>xpression data), an R package which provides a thorough exploration of both individual and combined gene expression, supplemented by pathway-guided analyses. VIBE’s versatility proves pivotal for disease stratification and therapeutic targeting in cancer, immune, metabolic, and other disorders.

**Results:** VIBE offers a wide array of functions that streamline the visualization and analysis of transcriptomics data for single- and dual-targeting therapies such as antibodies. Its intuitive interface allows users to evaluate the expression of target genes and their associated pathways across various indications, aiding in target and disease prioritization. Metadata, such as specific treatment or number of prior lines of therapy, can be easily incorporated to refine the identification of patient cohorts hypothesized to derive benefit from a given drug. Through real-world scenario representations using simulated data, we demonstrate how VIBE can be used to assist in indication selection for several user cases. VIBE integrates statistics in all graphics, enabling data-informed decision-making. Its enhanced user experience features include boxplot sorting and group genes either individually or averaged based on pathways, ensuring custom visuals for insightful decisions. For a deeper dive into its extensive functionalities, please review the vignettes on the GitHub repository (https://github.com/genmab/VIBE).

**Conclusions:** VIBE facilitates detailed visualization of individual and cohort-level summaries such as concordant or discordant expression of two genes or pathways. Such analyses can help to prioritize disease indications that are amenable to treatment strategies like bispecific antibody therapies or pathway-guided monoclonal antibody therapies. By using this tool, researchers can enhance the indication selection and potentially accelerate the development of novel targeted therapies with the end goal of precision, personalization, and ensuring treatments align perfectly with individual patient needs across a spectrum of medical domains.

## Background

The success of many therapeutic strategies hinges on the accurate targeting of pathological sites, specific cell types or mechanisms within the body^1^. Target expression plays a pivotal role in this context, ensuring that the therapeutic agents precisely address the malfunctioning cells or proteins, thereby minimizing collateral damage to healthy tissues^2^. A prime example of such precision targeting is found in immunotherapy where antibodies stand out due to their intrinsic ability to bind with high specificity to their target antigens. Antibody therapy, leveraging the inherent structure and specificity of antibodies, provides a precise method for treating a range of diseases including cancer, rheumatoid arthritis, Crohn’s disease, asthma, and many others^3-13^. Antibodies, composed of two antigen-specific Fab regions and an Fc region interacting with immune components, facilitate diverse therapeutic effects based on either agonizing or antagonizing a target, such as direct tumor growth inhibition, apoptosis induction, and/or recruitment or inhibition of immune-mediated mechanisms. Genetic engineering broadens these functionalities, for example by enabling dual-targeting antibodies that bind two distinct antigens, either on the same or different cells. One example of this strategy is the therapy Epcoritamab (DuoBody®-CD3xCD20), a bispecific antibody recently approved for diffuse large B-cell lymphoma, which binds CD20 on tumor cells and CD3 on T cells, enhancing T cell-mediated tumor kill^14-17^.

In addition to the target antigens themselves, a multitude of other genes – particularly those involved in downstream signaling pathways of the target gene(s) – may significantly influence the therapeutic efficacy^18^. Evaluating the mRNA expression of these genes to identify indications with high antigen expression, is one method to improve the selection of promising drug candidates for further study. Further, mRNA co-expression between transcripts can elucidate the underlying molecular mechanisms and potential resistance pathways, thereby informing the design of more effective combination strategies^19-22^.

While many existing tools allow for visualization of individual gene-level expression^23-25^, currently no tool exists that offers a comprehensive statistical visualization tailored for pathway-guided single- and dual-targeting antibodies. This functionality is critical to understanding inter-gene relationships and pathway dynamics, which can unlock the potential of targeted antibody therapies.

Here, we present VIBE, an R package uniquely tailored for the <u>Vi</u>sualization of <u>B</u>ulk RNA <u>E</u>xpression data applicable to technologies such as bulk RNA-seq, EdgeSeq, and microarray technologies. With VIBE, researchers can delve into the expression of specific genes or gene pairs within large transcriptomic datasets, such as the publicly available TCGA^26^, GTEx^27, 28^ or XENA^29^ databases, as well as custom datasets. Moreover, VIBE provides averaged scores for gene sets, such as pathway-associated genes, informing a comprehensive overview of gene interactions and functions within the pathway of interest. The resultant data can be visualized with boxplots and heatmaps, for an accessible and informative method to explore and interpret complex genomic data.

## Implementation

VIBE is relevant for R programming users across various skill levels; beginners can use the package’s base functions for easy data visualization while the package provides flexible control for proficient R programmers and the ability for experts to customize and extend the existing codebase. The package offers a wide range of functions, each with customizable parameters, although not all aspects are covered in detail here. Users can refer to the vignette available on the GitHub page. A dummy data set is included upon installation to enable out-of-the-box exploration of VIBE’s functionality. VIBE focuses on visualizing differences in gene/pathway expression patterns across user-defined patient groups.

### Generating simulated data set

The simulated data was generated using the rnorm^19^ package with mean and standard deviation for 35 genes (including “Tumor target” and “Immune target”) for 16 solid tumor types (**Table 1**) pre- and post-treatment. The Stringi^19^ package was used to randomly generate the patient IDs and sample IDs. The patient IDs were generated to match pre- and post-treatment data. For every tumor type, the “dummy” log2 TPM values were randomly generated for 100 patients, 50 each for pre- and post-treatment samples (Table 1). Users can define categories based on their research question. To illustrate this process, an additional parameter was added such that patients were assigned to two different databases (database1 or database2).

**Table 1.** Description of VIBE’s simulated dummy dataset.

| Indication | Abbreviated indication | Paired pre- and post-treatment samples | Unmatched database samples |  |
| --- | --- | --- | --- | --- |
|  |  |  | database1 | database2 |
| Bladder cancer | BLCA | 50 | 52 | 48 |
| Breast cancer | BRCA | 50 | 100 | 0 |
| Cervical cancer | CERV | 50 | 50 | 50 |
| Colorectal cancer | CRC | 50 | 62 | 38 |
| Head and neck squamous cell Carcinoma | HNSCC | 50 | 50 | 50 |
| Neuroendocrine tumors | NET | 50 | 50 | 50 |
| Non-small lung cancer | NSCLC | 50 | 0 | 100 |
| Ovarian cancer | OV | 50 | 56 | 44 |
| Pancreatic adenocarcinoma | PDAC | 50 | 36 | 64 |
| Prostate cancer | PRAD | 50 | 44 | 56 |
| Renal cell carcinoma | RCC | 50 | 38 | 62 |
| Sarcoma | SARC | 50 | 54 | 46 |
| Small cell lung cancer | SCLC | 50 | 50 | 50 |
| Stomach adenocarcinoma | STAD | 50 | 64 | 36 |
| Triple-negative breast cancer | TNBC | 50 | 50 | 50 |
| Uterine carcinoma | UTEN | 50 | 54 | 46 |

### Harmonizing the dummy dataset

The dataset input for VIBE should consist of the following essential columns: i) patient ID, ii) sample ID, iii) gene or feature name, iv) expression values, v) the unit used for plotting captions, and vi) grouping or plotting columns such as indication and treatment. To ensure the dataset is structured optimally for analysis and visualization, VIBE offers the *harmonize_df()* function. This function harmonizes the dataset by updating column names and generates additional columns that can serve as grouping variables for statistical analysis or visualization purposes (**Figure 1A**). Users have the flexibility to choose which additional columns to retain, enabling them to create multiple comparative visualization schemes tailored to their specific research needs.

**Figure 1.**
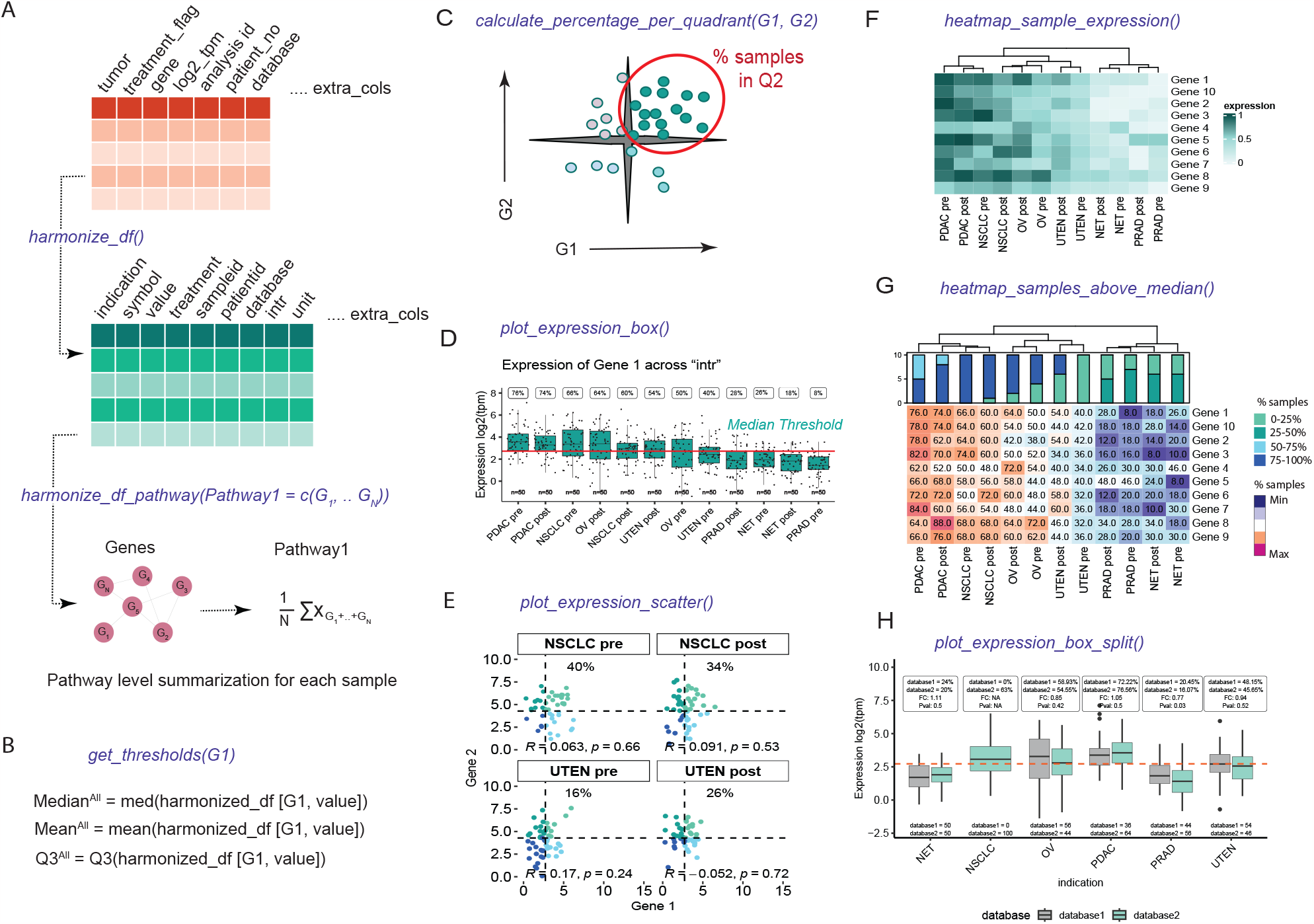
Implementation and overview of the statistics and visualization capabilities of the VIBE package. (A) The *df_harmonize()* function harmonizes dummy data, creating a structured dataframe with appropriate column names and format for VIBE functionalities. The *harmonize_df_pathway()* function combines genes into pathways for VIBE visuals. (B) The *get_thresholds()* function defines gene-specific thresholds (mean, median or 75% quantile (Q3)) for expression analysis and indication selection. (C) The *calculate_percentage_per_quadrant()* function extends *get_thresholds()* for dual-gene analysis, displaying percentage of samples in quadrants, correlation, and statistical significance (Spearman correlation). (D) *plot_expression_box()* presents a boxplot with sample distribution, thresholds, percentage satisfying thresholds, and user-defined grouping variables and groups (i.e. “intr”) generated when harmonizing the dataframe. (E) *plot_expression_scatter()* plots a scatterplot with user-selected grouping variables and groups and statistics calculated by *calculate_percentage_per_quadrant()* function. (F) *heatmap_sample_expression()* visualizes gene expression via heatmaps, allowing grouping variable selection, group plots, gene splitting, and ordering of genes. (G) *heatmap_samples_ above_median()* represent the percentage of samples satisfying threshold for all the genes with similar functionalities as mentioned in *heatmap_sample_expression()*. (H) *plot_expression_box_split()* function visualizes the expression of Gene 1 grouped by additional grouping columns with statistics (% above median for each group, the fold change (FC) and Kruskal-Wallis p-value (KW p)) above.

In line with the pathway-supported decision-making process, VIBE provides the *harmonize_df_ pathway()* function. This function offers users the capability to define a list of genes representing gene signatures or pathways. Utilizing this information, the function calculates the average expression values of all the genes within each gene signature or pathway. The *harmonize_df_pathway()* function then returns a structured data frame that is ready for further visualization using VIBE’s functionalities.

#### Defining thresholds

Rather than imposing arbitrary thresholds for identifying indications or classifying samples into high or low expression bins, our methodology allows users to pre-define thresholds based on the mean, median, or 75% quantile. This is achieved using the *get_threshold()* function in VIBE (**Figure 1B**). For consistency throughout this manuscript, we have chosen the median as the threshold.

To calculate thresholds, the median mRNA expression of each selected gene was computed separately for all categories (Median^category^). Subsequently, the median expression of all samples (Median^All^) was determined to dichotomize the category binarily. Specifically, if Median^category^ was greater than Median^All^, the category is considered to be of interest for the new therapeutic; otherwise, it is not. Additionally, the percentage of samples per category per dataset was calculated where Median^category^ is greater than Median^All^. This approach allows for a more flexible and user-defined thresholding strategy, enabling meaningful analysis and comparisons across different datasets and indications.

### Analysis of gene associations and gene-pathway interactions

One of the distinctive features that set VIBE apart is its capacity to visualize complex interactions and correlations between multiple genes or pathways simultaneously. For composite assessments of 2 genes or pathways, VIBE utilizes the previously generated thresholds to classify samples into 4 quadrants based on the expression levels of the selected genes or pathways. Users have the flexibility to choose any quadrant for visualization and can use the *calculate_percentage_per_ quadrant()* function to calculate the percentage of samples within the chosen quadrant (**Figure 1C**). Additionally, VIBE calculates the Spearman correlation between the selected genes or pathways, providing insights into their potential interactions and associations.

For composite assessment of antibodies versus multiple genes or pathways, VIBE calculates the percentage of samples within the chosen quadrant for each comparison. These results can be effectively visualized in a heatmap, allowing users to easily discern and interpret patterns and trends across various comparisons.

### Statistical analysis and visualization

The visualization capabilities of the VIBE package extend beyond basic graphics, enabling researchers to draw meaningful statistical inferences directly from the plots. The visuals are extended using ggplot2^19^ and ComplexHeatmap^19^ functionalities. The *plot_expression_box()* function, for instance, provides a boxplot that incorporates the threshold and percentage of samples satisfying the threshold within the plot itself. This eliminates the need for a separate table and facilitates the selection of user-defined groups (**Figure 1D**). Additionally, the *plot_expression_scatter()* function offers a scatterplot between two genes, where quadrant-specific colors are defined by the thresholds of the two genes. This scatterplot not only visualizes the percentage of samples in the selected quadrant but also displays the correlation between the two genes and its statistical significance within the given dataset using Spearman correlation (**Figure 1E**). The functions provided also enable the plotting of specific indication categories. To ensure statistical robustness, the thresholds for these categories are calculated using the complete dataset. Visualizations are then generated specifically for the categories chosen by the user.

For a comprehensive overview of relevant genes in the dataset, VIBE offers the heatmap representation of gene expression in user-defined groups (**Figure 1F**). Moreover, the heatmap also shows the percentage of samples that are above the user-defined threshold, aiding researchers in identifying significant gene expression patterns (**Figure 1G**). Furthermore, the *plot_expression_box_ split()* function accommodates additional grouping variables, as an extension of *plot_expression_ box()* with statistics (% above median for each group, the fold change (FC) and Kruskal-Wallis p-value (KW p)) depicted above . This function allows users to compare the expression of a gene in control versus treatment conditions, providing valuable insights into the impact of different conditions on gene expression.

VIBE offers additional advanced visualization and statistical inferences for comprehensive genomic data analysis. Demonstrated in its vignette and results section, VIBE enables researchers to identify indications of interest, assess gene and pathway correlations, and make data-driven decisions, enhancing precision in drug development and disease research.

## Results

The robust capabilities of VIBE were demonstrated through two compelling case studies. In the first case study, the statistical and visual representations were leveraged to effectively identify cancer indications of interest for a novel antibody-drug conjugate (ADC). In the second case study, VIBE is shown how to empower researchers to make informed decisions beyond target expression. With VIBE’s comprehensive pathway and gene signature analyses, researchers can assess the potential impact of multiple pathways in tandem with target expression. This holistic approach provides valuable support in making well-informed decisions regarding therapeutic interventions, enhancing the precision and effectiveness of treatment strategies.

### VIBE as a tool for identifying indications of interest for a novel antibody-drug conjugate (Case study 1)

In this case study, VIBE identified potential indications of interest for a novel ADC directed against the ‘Tumor target’ gene. In the dummy data, high expression levels of the “Tumor target” gene are a relevant hypothesis that may reflect high efficacy for this targeting strategy. The expression levels of the target are visualized using boxplots across the cancer types which revealed that ‘Tumor target’ gene expression is the highest in CRC, SCLC, PRAD, UTEN, NET and STAD, suggesting these as potential candidates for the ADC therapy (**Figure 2A**). However, these findings should still be confirmed using single cell approaches to determine whether the “Tumor target” gene is indeed highly expressed by cancer cells, and not healthy (immune) cells. In addition, protein expression in the top indications should be confirmed in further experiments, for example by immunohistochemistry.

**Figure 2.**
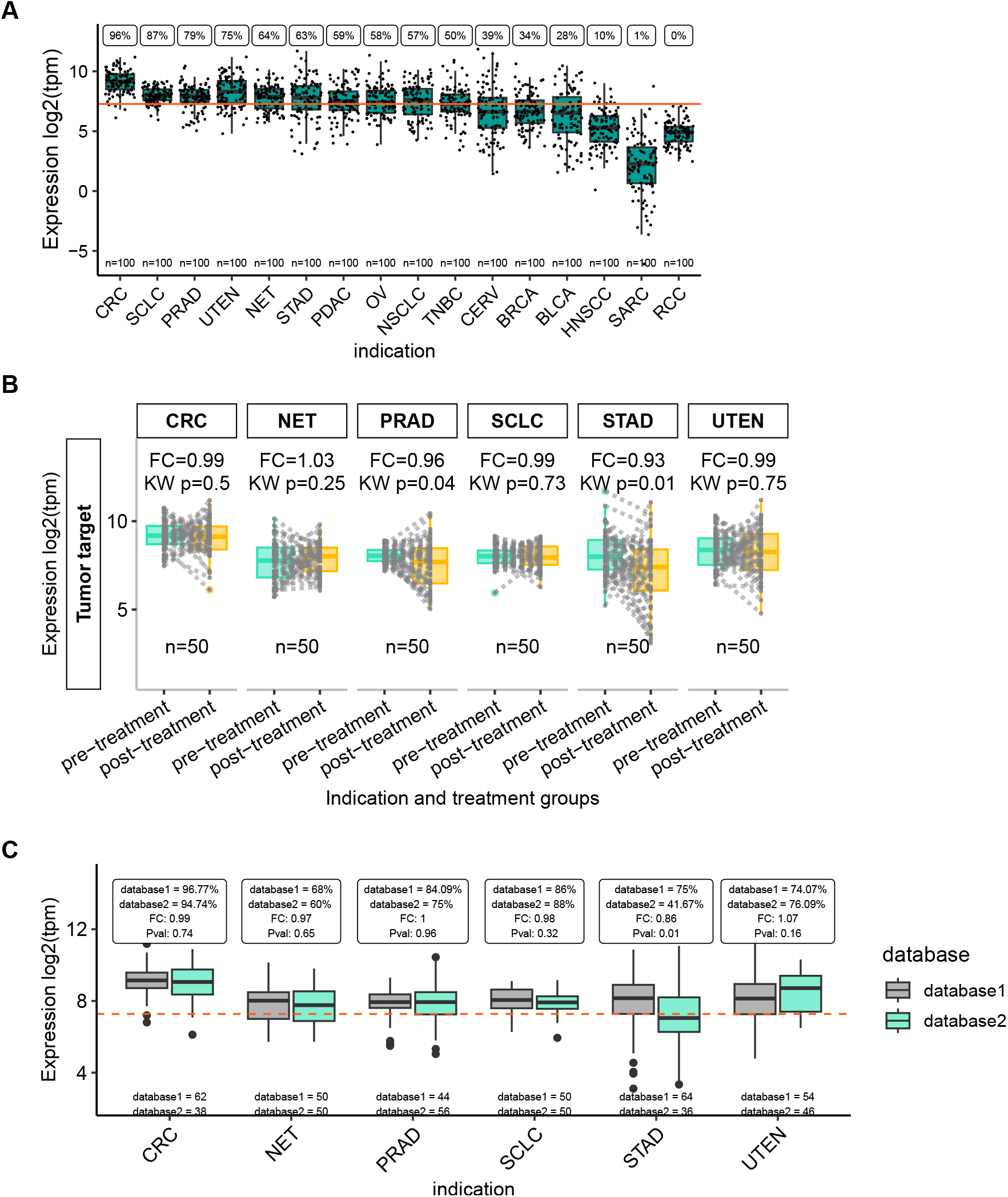
“Tumor target” expression highlights indications of interest for a novel ADC. (A) VIBE’s *plot_expression_ box* function shows a box for each group of interest, indication in this example, showing simulated data included in the VIBE package. Dots indicates individual samples; the orange line indicates the median expression of the plotted gene across all samples in the dataset. The percentage of samples with higher than median expression is printed above each box, while the number of samples in each group is shown just above the x-axis. (B) The *plot_box_pre_post* function plots a box for pre- and post-treatment samples, with each dot representing a sample, while lines connect paired samples. The fold change (FC) and Kruskal-Wallis p-value (KW p) are printed above each group of interest. (C) The *plot_expression_box_ split* function allows for the comparison of non-matched data and includes the fold change (FC) and Kruskal-Wallis p-value (Pval) between the two groups. The number of samples included in each dataset is printed just above the x-axis. The orange dotted-line indicates the median expression for this gene across all samples in the combined dataset. Indication abbreviations are explained in Table 1

As clinical trials for a new cancer drug are often performed in a population of patients who have already received other treatment(s), it is important to study the difference between pre-and post-treatment samples when it comes to expression of the drug’s target. VIBE’s specialized function *plot_ box_pre_post()* plots paired pre- and post-treatment samples and calculates the fold-change and p-value between the two states. Using this functionality in the dummy dataset, STAD was identified as the tumor type where the “Tumor target” expression was lower in post-treatment samples (**Figure 2B**). This indicated that the ADC therapy could be less effective in post-treatment STAD patients compared to pre-treatment patients.

The users would like to include additional labels or groups of interest (e.g. unmatched samples) in their RNA-seq dataset. To exemplify such an analysis, the dummy data was harmonized while keeping the additional column representing unmatched samples from “database1” and “database2” (**Figure 1A**). VIBE’s function *plot_expression_box_split()* allows comparison of gene expression differences between the unmatched samples. In the dummy data, the “Tumor target” expression was lower in STAD samples in database 2 compared to database 1 (**Figure 2C**), indicating that the underlying data for this indication warrants further investigation. Such a functionality of VIBE allows comparing tumor to normal samples e.g. GTEx vs TCGA in a real-world setting.

### VIBE as a tool for pathway-guided indication selection for a novel mono- or bi-specific antibody (Case Study 2)

In this case study, the analysis involving more than two genes or pathways was conducted. Depending on the intended mechanism of action (MoA), the efficacy of a novel therapy may depend on the presence of additional genes or even entire pathways. Consequently, high expression of specific genes and pathways may inform hypotheses of improved efficacy for the therapy under consideration. As an example, this case study aimed to select cancer indications for a bi-specific antibody targeting both tumor (“Tumor target”) and immune (“Immune target”) cells. This bi-specific antibody’s mechanism of action (MoA) promotes immune cell-tumor cell interaction, leading to immune cell activation, and resulting in tumor cell kill by the immune cell which is visualized here along with the two targets.

To exemplify the capabilities of VIBE, we use the “Immune target” and “Tumor target” variables from the dummy dataset to visualize targets from a bispecific antibody therapy. Only the indications with median expression higher than the threshold for the “Tumor target” (Case study 1) were selected for further analysis. It’s important to note that these threshold values continue to be calculated based on the entire dataset to ensure statistical validity and robustness. By examining the percentage of samples falling within the second quadrant of the scatterplot (**Figure 3A**) and the corresponding bar plot (**Figure 3B**), we identify PDAC pre-treatment, STAD pre-treatment, CRC pre-treatment, and CRC post-treatment as potential indications and treatment groups of interest for the bispecific antibody.

**Figure 3.**
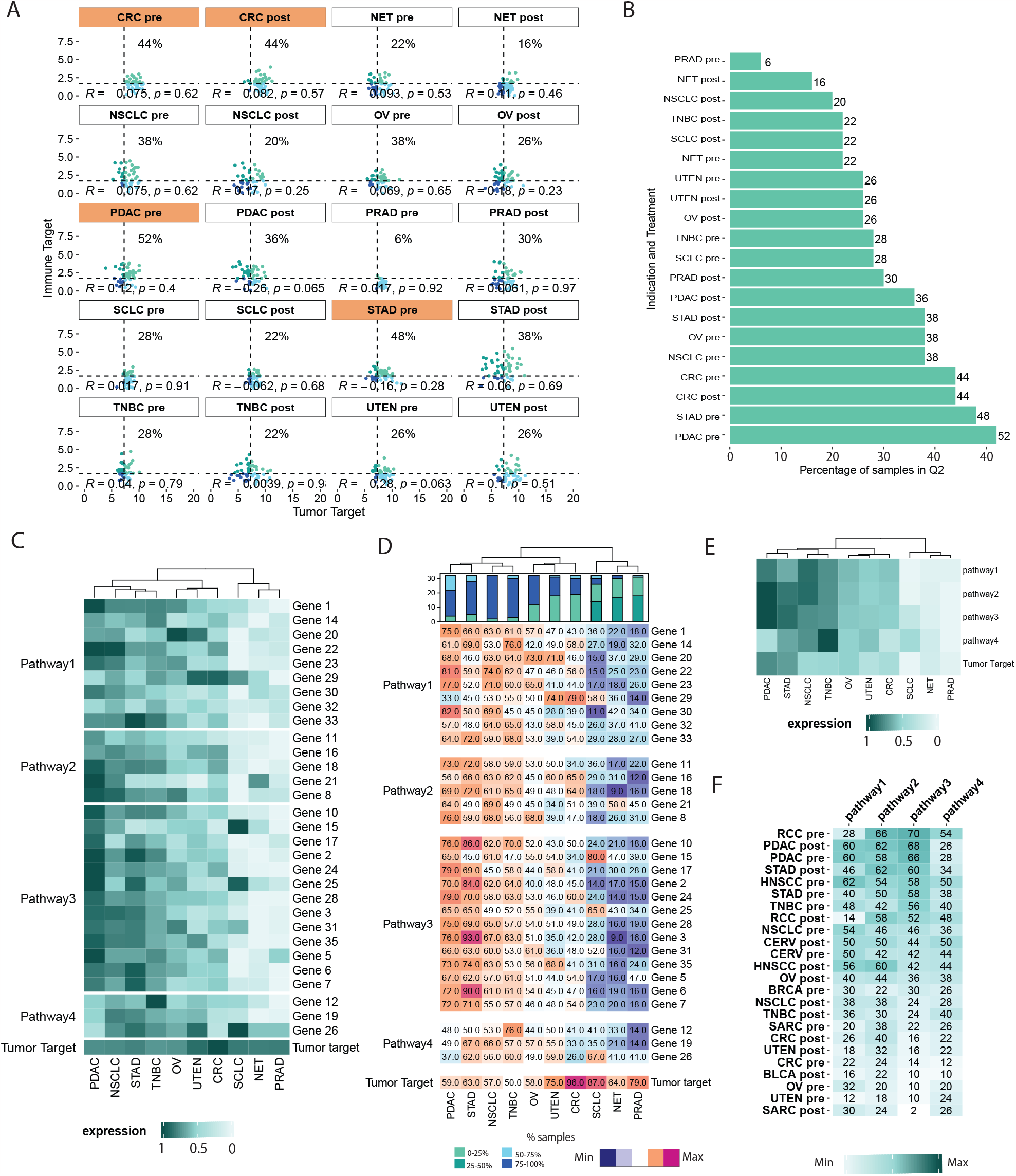
Identification of indications for bi-specific or pathway-guided single ADC. (A) Scatterplot representing the distribution of samples in 4 different quadrants based on the user-defined threshold (“Tumor target” vs “Immune target”). VIBE’s *plot_expression_scatter()* function generates a FACS-like scatterplot, dividing samples into four quadrants based on the median expression of “Tumor target” and “Immune target” across all samples. The *plot_groups* argument of the function was used to select the indications as shown in Figure 1A. Indications and treatment groups with over 40% of samples expressing high levels of both “Tumor target” and “Immune target” are highlighted in orange. (B) Boxplot representing the percentage of samples in user-defined groups and quadrants. The *plot_perc_pop()* function in VIBE generates a boxplot that displays the percentage of samples expressing high levels of both “Tumor target” and “Immune target”. (C) Heatmap representing the expression of user-defined grouping of genes (pathways) in user-defined/selected indications. VIBE’s heatmap_sample_expression() function is based on the ComplexHeatmap package^19^, enabling easy grouping of genes into pathways and visualization of their expression along with the target of interest, such as the “Tumor Target” in this case. One can easily visualize the change in the expression of genes in pathways compared to the “Tumor target”. (D) Heatmap representing the percentage of samples in user-defined grouping of genes (pathways) and user-defined/selected indications. VIBE’s *heatmap_samples_above_median()* function, also based on ComplexHeatmap^19^, allows users to visualize the percentage of samples with high expression levels of individual genes within the defined pathways. (E) Heatmap representing the expression of pathways in user-defined/selected indications. VIBE’s *heatmap_sample_expression()* function allows users to visualize the average expression of genes in pathways in conjunction with the “Tumor Target”. Comparing the composite pathway expression to individual genes, as shown in Figure 2C, offers a more robust approach for making decisions on indication selection, particularly when considering the mode of the ADC’s mechanism. (F) Composite analysis of pathways vs ADC. VIBE’s *heatmap_composite_scores()* function generates a heatmap representing the percentage of samples in the Q2 quadrant of the scatterplot. The function allows the user to define a threshold to select indications based on the scores. In this case, a threshold of 20 was used.

The evaluation of pathways is crucial in assessing the effectiveness of antibodies with an MoA that depends on specific pathways and plays a pivotal role in the selection of suitable indications and treatment groups. For instance, in scenarios where mRNA signatures such as T-cell infiltration or activation play a vital role, high expression of genes in these pathways is essential to represent an inflamed, or “hot tumor” scenario for the intended MoA to become a reality. VIBE offers two distinct scenarios for pathway assessment: (1) evaluating individual genes within a pathway, and (2) composite analysis of pathways using summarized expression of signature genes. To represent the first scenario, heatmap visualizations can be used to illustrate gene expression (**Figure 3C**) and the percentage of samples with expression above the median threshold (**Figure 3D**). In the second scenario, an averaged expression of genes to represent pathways, alongside the “Tumor target” in a heatmap (**Figure 3E**). Using these visuals, PDAC and STAD emerged as relevant indications for further investigation.

Although scatterplots, as shown in Figure 2A, offer detailed insights into quadrant-specific sample distributions, they become complex when comparing one target with multiple genes or pathways. To address this, the heatmap_composite_score() function presents a user-defined quadrant-specific percentage of samples in a heatmap (**Figure 3F**). This allows users to quickly assess and select indications of interest. For example, NSCLC pre-treatment and PDAC pre-treatment patient cohorts show high expression of both the “Tumor target” and Pathways 1, 2, and 3, (e.g. T-cell activation, T-cell exhaustion or T-cell infiltration) making them potentially relevant indications. On the other hand, SCLC pre- and post-treatment patient cohorts would be of interest, showcasing high expression of the “Tumor target” and pathway 4 (e.g. NK cell signature). Users can flexibly choose the pathways and targets to be represented in the analysis.

This comprehensive approach allows researchers to gain a deeper understanding of the interactions and functional significance of gene pathways, facilitating informed indication selection for therapeutic antibody development. In summary, VIBE’s comprehensive visualization capabilities enable researchers to explore and compare complex interactions between the drug’s target gene(s) and multiple genes or pathways, aiding effectively in the selection of potential indications for therapy development.

## Applications

The VIBE package is a unique visualization tool for developing hypotheses around disease stratification and targeted therapy. This tool facilitates composite analyses of multiple genes or pathways with user-defined thresholds, enabling researchers to understand gene expression dynamics comprehensively. This is instrumental for making informed and data-driven decisions in antibody development and targeted therapeutic design.

Especially pertinent as antibody therapies prove effective against diseases like cancer and auto-immune disorders, VIBE aids in indication selection. It is also useful in dissecting drug resistance mechanisms by highlighting gene expression and pathway alterations associated with drug resistance to develop strategies for overcoming treatment challenges. Moreover, its ability to represent intricate genes and pathways interactions makes it valuable for systems biology research by providing insights into broader regulatory networks and systems. In essence, VIBE offers insights into gene expression dynamics across vital pathways, with broad implications for advancing personalized treatments and understanding disease intricacies.

## Conclusions

In our evaluation, the VIBE package stands out for its statistics-driven approach to data visualization of gene expression data. This tool not only integrates statistics in visualisations for single-gene exploration but also provides comprehensive visualization tailored for pathway-guided single- and dual-targeting antibodies. The visuals including expression levels of gene(s), genes corresponding to the related MoAand the statistics are instrumental in making data driven decisions to the indication identification process. Unlike existing tools, VIBE offers a statistics-centric visualization approach, granting users the flexibility to work across various datasets and disease models.

## Availability and requirements

Project name: VIBE

Project home page: https://github.com/genmab/VIBE

Operating system(s): Platform independent

Programming language: R

Other requirements: none

License: MIT license

Any restrictions to use by non-academics: None

### List of abbreviations

Antibody-Drug: Combination:
ADC: Bladder Cancer:
BLCA: Breast Cancer:
BRCA: Cervical Cancer:
CERV: Colorectal Cancer:
CRC: Genotype-Tissue Expression:
GTEx: Head and Neck Squamous Cell Carcinoma:
HNSCC: Mechanism of Action:
MoA: NeuroEndocrine Tumors:
NET: Non-Small Cell Lung Cancer:
NSCLC: Ovarian Cancer:
OV: Pancreatic Adenocarcinoma:
PDAC: Prostate carcinoma:
PRAD: Renal Cell Carcinoma:
RCC: Sarcoma:
SARC: Small Cell Lung Cancer:
SCLC: Stomach Adenocarcinoma:
STAD: Triple-Negative Breast Cancer:
TNBC: VIzualization of Bulk RNA Expression data:
VIBE: The Cancer Genome Atlas:
TCGA: Uterine Carcinoma: UTEN.

## Declarations

### Ethics approval and consent to participate

Not applicable

### Consent for publication

Not applicable

### Availability of data and materials

The simulated data used in this study are included as party of the VIBE R package. https://github.com/genmab/VIBE/data

## Authors’ contributions

ICRM conceptualized the study. IK, SDvA, ICRM and FB carried out the programming and design of the package. LFM Tested the package. BH and CK contributed to scientific discussions of the package. All authors contributed to the writing and/or editing of the manuscript.

## Acknowledgements

A special acknowledgement to Francis Blokzijl whose original work laid the groundwork for the development of this R package. The Scientific Communication and Scientific publication team at Genmab for their input on the manuscript.

